# Frontostriatal Functional Connectivity Underlies Self-Enhancement During Social Evaluation

**DOI:** 10.1101/2021.05.07.443090

**Authors:** Michael H. Parrish, Janine M. Dutcher, Keely A. Muscatell, Tristen K. Inagaki, Mona Moieni, Michael R. Irwin, Naomi I. Eisenberger

## Abstract

Self-enhancement, the tendency to view oneself positively, is a pervasive social motive widely investigated in social and personality psychology. Despite research on the topic over the past several decades, relatively little is known about the neurocognitive mechanisms underlying this motive, specifically in social evaluative situations. To investigate whether positive emotion regulation circuitry, neural circuitry involved in modulating positive affect, relates to the operation of the self-enhancement motive in social contexts, we conducted an fMRI study in a sample of healthy young adults. We hypothesized that self-enhancement indices (state, trait self-esteem) relate to greater positive functional connectivity between right ventrolateral prefrontal cortex (RVLPFC), a neural region implicated in emotion regulation, and the ventral striatum (VS), a region associated with reward-related affect, during a social feedback task. Following social-evaluative feedback, participants maintained stable self-esteem or experienced a drop in state self-esteem. We found stable state self-esteem and higher trait self-esteem related to greater positive connectivity in RVLFPC-VS circuitry during receipt of positive (vs. neutral) feedback. These findings implicate the neurocognitive mechanisms underlying emotion regulation in the functioning of the self-enhancement motive and highlight a pathway through which self-enhancement may restore feelings of self-worth during threatening circumstances.

---

Self-enhancement, the tendency to view oneself in a favorable manner, is a core human social motive and is found, in some form, within all cultures (Fiske, 2019; Sedikides & Gregg, 2008). Across several countries, individuals have been shown to self-enhance on personality traits that are deemed personally-relevant to themselves or to their cultures more broadly (Sedikides et al., 2003). In fact, the universality and primacy of self-enhancement has been confirmed by a large-scale meta-analysis using over 500 diverse, independent samples (Mezulis et al., 2004). Indeed, self-enhancement may be a pervasive part of human nature because enhancing personal attributes or values has evolved as a useful cognitive adaptation to protect one’s self-view from the challenging or stressful events we face on a daily basis (Alicke, 1985; Shelley E. Taylor & Sherman, 2008). Attesting to its importance, research into self-enhancement as a psychological coping strategy has been one of the most widely studied topics in social and personality psychology and has been a mainstay in psychology since the days of Freud (Alicke & Sedikides, 2011). However, despite the steady stream of research on the topic of self-enhancement in social and personality psychology to date, research has yet to reach a consensus about the psychological or neural mechanisms underlying self-enhancement. The current study focuses on using neuroimaging to explore whether emotion regulation circuitry is critical for self-enhancement.

Behavioral studies in social psychology suggest a strong link between self-enhancement related processes and emotion regulation mechanisms. These studies point to an important role for self-enhancing strategies in increasing positive affect in particular. In fact, some have even speculated that mood regulation and the restoration of positive affect is the primary functional benefit of self-enhancement (Sedikides & Gregg, 2008; Tesser, 2000). For example, behavioral research suggests that individuals with positive self-views engage in regulatory processes in order to “savor” or enhance positive feelings in the context of self-relevant events (Bryant, 1989, 2003; Wood et al., 2003, 2005). Specifically, following the description of a positive and personally relevant event, individuals with greater self-enhancing tendencies tended to enjoy and magnify their success (Wood et al., 2003), whereas individuals with less self-enhancing tendencies tended to mute or dampen the positive feelings associated with the event. In addition, related research shows that self-serving judgments about the desirability or favorability of one’s own qualities are used strategically to regulate affective experience in order to improve negative mood (Roese & Olson, 2016). For example, in response to failures, self-enhancers often seek out opportunities to repair their mood by increasing positive affect (e.g., watching comedic videos) and improve self-related feelings (e.g., reinterpreting feedback in a self-serving manner; Alicke & Sedikides, 2009; Heimpel et al., 2002). Relatedly, self-enhancers tend to report overly optimistic attitudes about others’ future perceptions of their own behaviors when facing the threat of social evaluation (Preuss & Alicke, 2009). Moreover, individuals with self-enhancing tendencies often maintain an inflated self-image in the face of adversity through the use of positive affirmations of their personal values (Taylor & Armor, 1996; Taylor & Sherman, 2008). Taken together, these studies suggest that the mechanisms of self-enhancement are most likely rooted in the mechanisms of domain-general positive emotion regulation, specifically regulatory mechanisms that increase positive affect.

Neuroscience approaches can be utilized to investigate whether emotion regulation-related neural mechanisms underlie individual differences in self-enhancement. With regard to the specific regulatory mechanisms that might account for self-enhancement effects, both upregulation and downregulation of positive affect has been tied to ventral striatum functioning (Greening et al., 2014; Kim & Hamann, 2007; Wager et al., 2008). More specifically, connectivity between the ventrolateral prefrontal cortex (VLPFC) and the ventral striatum (VS) may be an important regulatory circuit implicated in positive reappraisal (Wager et al., 2008). Research has shown that the effect of right VLPFC (RVLPFC) activation on emotion regulation outcomes is mediated in part by reward-related VS activation (Wager et al., 2008). Taken together, previous research in psychology and neuroscience points to an important role for positive emotion regulation circuitry, specifically those circuits involved in modulating reward-related activity, in maintaining favorable self-views and the motivational processes which underlie them.

To date, however, few studies have focused on the neural underpinnings of self-enhancement effects and the ones that have, have mostly focused on self-referential processes rather than positive emotion regulation mechanisms (Beer & Hughes, 2011; Chavez & Heatherton, 2015; Hughes & Beer, 2012a, 2012b). Specifically, in most earlier neuroimaging studies examining self-enhancement, participants were instructed to rate the self-relatedness of a personality trait adjective or to explicitly compare themselves to their peers (Beer, 2014; Beer & Hughes, 2010). Although this is a well-known method for assessing the presence of self-enhancement, this task may be less similar to the real-life situations in which self-enhancement processes are engaged, such as in response to evaluative feedback, and thus may be less likely to recruit emotion regulatory processes. This may be one of the multiple reasons why previous investigations have yet to find reliable associations between self-enhancement effects and activity in either reward-related regions or positive emotion regulation circuitry (Beer & Hughes, 2011; Chavez et al., 2017; Flagan et al., 2017; Izuma et al., 2018; Kuzmanovic et al., 2016). Thus, neuroimaging research employing naturalistic tasks which simulate real-life situations in which self-enhancing processes are utilized, such as in response to social evaluative feedback, is critically needed.

In order to examine a possible circuit-level neural correlate of self-enhancement, we administered an experimental fMRI task in which subjects would have a chance to self-enhance in response to receiving positive and negative feedback from others. Participants completed an fMRI scan in which they received feedback about how an evaluator ostensibly rated their personality and opinions. A third of the feedback they received was positive, a third was neutral, and a third was negative. They were told that this feedback came from a peer (who was actually a confederate) who listened to a previously-recorded audio interview in which the participant discussed themselves. Prior to the scan, participants completed standard questionnaire measures of trait self-esteem. Before and after the evaluative task in the scanner, participants additionally indicated their state self-esteem, and we calculated whether subjects showed a drop in their state self-esteem from pre-to post-task, or whether their state self-esteem remained stable (did not change). Based on prior work (Eisenberger et al., 2011) and since subjects received considerable negative feedback during the task, we did not expect state self-esteem to increase. Individuals who maintained a stable level of state self-esteem in response to this evaluative task were inferred to have been more likely to have engaged in self-enhancement processes during the task itself compared to those who showed a drop in state self-esteem.

To examine whether emotion regulation circuitry plays a role in the act of self-enhancement, we examined whether RVLFPC-VS functional connectivity positively related to indices of self-enhancement (i.e., trait and state self-esteem). We hypothesized that stronger functional connectivity in the RVLPFC-VS circuit in response to positive and negative evaluative feedback would be related to stable levels of state self-esteem, as well as relatively higher levels of trait self-esteem.

## Methods

### Participants

Fifty participants (23 female; M= 23.36 years, range= 18-47 years) provided data for the present study. Participants were recruited from UCLA and the surrounding community. The study was generally representative of standard UCLA demographics: 48% White, 28% Asian/Pacific Islander, 16% Latino/Chicano, 2% Black/African American, and 6% Other. In total, one hundred fifteen participants were originally enrolled in a larger study (ClinicalTrials.gov identifier NCT01671150) for which this current study was a follow-up investigation. The purpose of the larger study was to examine the effects of inflammation on social and affective responses and thus half of the full sample of participants received an injection of an inflammatory challenge (0.8 ng/kg endotoxin; Moieni et al., 2015). All participants incorporated into the present follow-up investigation were a part of the placebo control group, and therefore did not receive the inflammatory drug. Neural responses to this task as a function of the inflammatory challenge have been reported previously (e.g., Muscatell et al., 2016). In this paper, we focus specifically on how functional connectivity related to individual differences in trait self-esteem and changes in state self-esteem from before to after the evaluative task in subjects in the placebo condition only.

### Procedure

Potential participants were excluded during a phone screening and an in-person screening due to contraindications for the MRI environment (e.g. metallic implants, left-handedness, claustrophobia) and a history of neurological or psychiatric disorders (through the SCID). They were also excluded if they had a history of physical health problems (e.g., allergies, autoimmune disease, BMI greater than 30, current prescription or recreational drug use). All participants provided written informed consent. The UCLA IRB approved all study procedures. Complete procedures for the study session have been explained in full elsewhere (Moieni et al., 2015). On the day of the experiment, participants arrived at the UCLA Clinical and Translational Research Center, where a nurse inserted a catheter into the dominant forearm for hourly blood draws and a catheter into the opposite forearm for continuous saline flush and drug (placebo) administration. Following administration of the drug/placebo, participants completed an audio-recorded interview during which they were asked about their personal characteristics and opinions for about 10 min (e.g., “What makes you happy?”, “What is your best quality?”, “What is your greatest shortcoming?”, and “What are you most afraid of?”). Participants were informed that, during the MRI scan, trained evaluators would listen to and form impressions of them based on their interview and that participants would rate how they felt in response.

### fMRI Social Feedback Task

Upon arrival at the MRI scanning center, participants met two other individuals (actually confederates; one male, one female) with whom they believed they would be interacting during the MRI tasks. Specifically, for the present task, participants were told that while they were in the MRI scanner, the evaluators would be seated in the scanner control room and would listen to the participant’s interview and that one of them would provide feedback about how the participant came across in the interview. In reality, participants in the scanner viewed the computer screen displaying an array of adjective “buttons” (i.e., “interesting,” “modest,” “boring”) and watched a pre-recorded video of a cursor moving around the screen, which they were led to believe was the real-time display of the confederate’s feedback on their interview.

The number of feedback adjectives selected were equally divided into a positive category (e.g. ‘intelligent’), a neutral category (e.g. ‘practical’) and a negative category (e.g. ‘annoying’). Participants watched as a new adjective button was selected every 10–12 s. During the entirety of the scan session, participants received fifteen each of positive, neutral, and negative feedback selections. All feedback was presented in a pseudorandom order with the constraint that no more than two adjectives of the same valence were presented consecutively. Following the experimental session, participants were promptly debriefed in a funneled manner and informed of the true purpose of the task. No participants reported suspicion prior to debriefing about the true purpose of the task.

### State Self-Esteem Measure

Participants rated their state self-esteem before starting the scan and again immediately after completing the scanning session. Based on previous studies (Leary et al., 1998), participants rated their state self-esteem by judging “how they felt right now” on a 4-point Likert scale from 1 (really bad) to 4 (really good). First, difference scores were calculated by subtracting pre-session state self-esteem from post-session state self-esteem. Then, participants were grouped based on whether they showed a decline in state self-esteem (n=19) or remained stable in state esteem (n=20) from before to after the social feedback categorized as either stable (no change) or declined (1-2 point change)). No participants showed an increase in state selfesteem.

### Trait Self-Esteem Measure

Trait self-esteem was measured approximately 2 hours prior to the scanning session using the Rosenberg Self-Esteem Scale (Rosenberg, 1965). This measure is the most widely used measure for global self-worth and has been shown to have good internal consistency and construct validity. This scale assesses self-views with items such as “I feel that I have a number of good qualities”. Ratings are on a 4-point Likert scale from 1 (strongly agree) to 4 (strongly disagree).

### MRI Data Collection

Imaging data were acquired from a Siemens 3 T Tim Trio MRI scanner at the UCLA Staglin IMHRO Center for Cognitive Neuroscience. A high-resolution T1-weighted echo-planar imaging volume (spin-echo, TR = 5000 ms; TE = 33 ms; matrix size 128 x 128; 36 axial slices; FOV = 20 cm; 3-mm thick, skip 1-mm) and T2-weighted, matched-bandwidth anatomical scan (slice thickness = 3 mm, gap = 1 mm, 36 slices, TR = 5000 ms, TE = 34 ms, flip angle = 90°, matrix size 128 x 128, FOV = 20 cm) were obtained for each participant. Afterwards, a functional scan was acquired which lasted 8 min, 38 s (echo planar T2-weighted gradient-echo, TR = 2000 ms, TE = 25 ms, flip angle = 90 °, matrix size 64 x 64, 36 axial slices, FOV = 20 cm; 3-mm thick, skip 1-mm).

### MRI Pre-processing

MRI data were pre-processed with the Statistical Parametric Mapping software (SPM8; Wellcome Department of Cognitive Neurology, London, UK). The pipeline for pre-processing incorporated functional realignment to correct for head movement, co-registration of the functional to the structural images, and spatial normalization of functional and structural images to Montreal Neurologic Institute (MNI) space (resampled at 3 mm isotropic), and spatial smoothing using an 8mm Gaussian kernel, full width at half maximum, to increase signal-to-noise ratio. The feedback task was modeled as a block design. The presentation of each feedback word (positive, negative, or neutral trait adjectives) and the subsequent 10 seconds were modeled as a block.

### Functional Connectivity Analyses

Functional connectivity analyses were conducted with the CONN toolbox (nitrc.org/projects/conn) implemented through MATLAB and SPM8 software. The preprocessed functional and structural data were entered into the toolbox. Confounding variables that distort functional connectivity values were removed through the CONN CompCor algorithm for physiological noise as well as temporal filtering (f>.008Hz). Realignment parameters (representing head movement) produced during pre-processing were also entered in the toolbox as nuisance covariates to be removed from statistical analyses. For the functional data collected during the social feedback task, condition onsets and durations (10 seconds for each block) were specified in the toolbox, so that BOLD time series could be appropriately divided into task-specific blocks.

For the main statistical tests of interest, we conducted ROI-to-ROI generalized psychophysiological interactions (gPPI) analyses to determine functional connectivity (i.e., temporal correlations) between the RVLPFC and both the left and right ventral striatum. For these analyses, we chose ROIs implicated in previous studies of emotion regulation (Frank et al., 2014). The RVLPFC ROI was generated by creating a spherical volume centered on peak coordinates in right inferior frontal gyrus (MNI 45, 22, 4) from a previous meta-analysis on the neural correlates of cognitive emotion regulation (Frank et al., 2014). The right and left ventral striatum ROIs were structurally defined using the AAL atlas (Tzourio-Mazoyer et al., 2002). The ventral parts of the right and left caudate nucleus and putamen from the atlas were constrained at x between 0 and −24, y between 4 and 18, and z between 0 and −12 for the left ROI and x between 0 and 24, y between 4 and 18, and z between 0 and −12 for the right ROI (based on ROIs from Inagaki & Eisenberger, 2012). Thus, we constrained the ROI to the ventral parts of the caudate nucleus and putamen to create this VS ROI.

Within the ROIs, the BOLD activation time series was averaged across all voxels. Functional connectivity (gPPI parameter estimates) values were computed on each individual’s feedback condition time series from these ROIs at the single-subject level. These connectivity values provide a measure of the statistical dependence of the ROIs’ BOLD activation time series. Connectivity values underwent Fisher’s *r-to-Z* transformation to ensure assumptions of normality. This procedure was completed to generate task-evoked absolute connectivity measures for each of the three social feedback conditions. Relative connectivity measures were generated by taking the difference between connectivity values produced by the positive trials vs. the neutral trials as well as by the negative trials vs. the neutral trials. In other words, they should be interpreted as the difference in the functional coupling between these neural regions (i.e., RVLPFC and VS) between these conditions. These gPPI parameter estimates values were imported into SPSS v23 for further statistical analyses. We first examined differences in RVLPFC-VS connectivity during positive (vs. neutral) feedback between the stable and decreased state self-esteem groups by performing independent samples *t*-tests as well as Mann-Whitley *U* tests due to concerns about data normality and outliers. We then performed the same analyses for RVLPFC-VS connectivity during negative (vs. neutral) feedback. Any significant effects were followed up by additional analyses exploring whether the effects were being specifically driven by absolute connectivity during the positive, negative, or neutral feedback condition. To examine correlations between trait self-esteem and RVLPFC-VS connectivity, we computed Pearson’s correlations and Spearman’s rank correlations between trait self-esteem and RVLPFC-VS connectivity during positive (vs. neutral) feedback as well as during negative (vs. neutral) feedback.

## Results

### RVLPFC-VS Functional Connectivity During Feedback Processing

#### Relationships with changes in state self-esteem

To examine whether subjects with stable state self-esteem were engaging in greater self-enhancement processes, we examined whether those who maintained stable state self-esteem in response to both positive and negative feedback showed greater RVLPFC-VS functional connectivity compared to those who showed a drop in state self-esteem.

First, with regard to neural activity to positive (vs. neutral) feedback, we found that the stable state self-esteem group showed significantly greater positive connectivity between RVLPFC and left VS (LVS) compared to the group that showed decreases in state self-esteem (*t*(37) = 2.948, *p* = .003; Fig. 1a). Moreover, as displayed in Figure 1, those whose state selfesteem stayed stable during the task showed positive functional connectivity on average, whereas those whose self-esteem decreased as a function of the task showed negative functional connectivity on average. Nonparametric statistical testing was additionally used due to concerns about data normality and outliers. A Mann-Whitley *U* Test also revealed the stable self-esteem group showed significantly greater positive connectivity, compared to the decreased self-esteem group (*U* = 278.00, *p* = .007). Results indicated that functional connectivity between the RVLPFC and the right VS was not significantly different between stable and decreased state self-esteem groups (*p* > .05).

**Figure 1.**
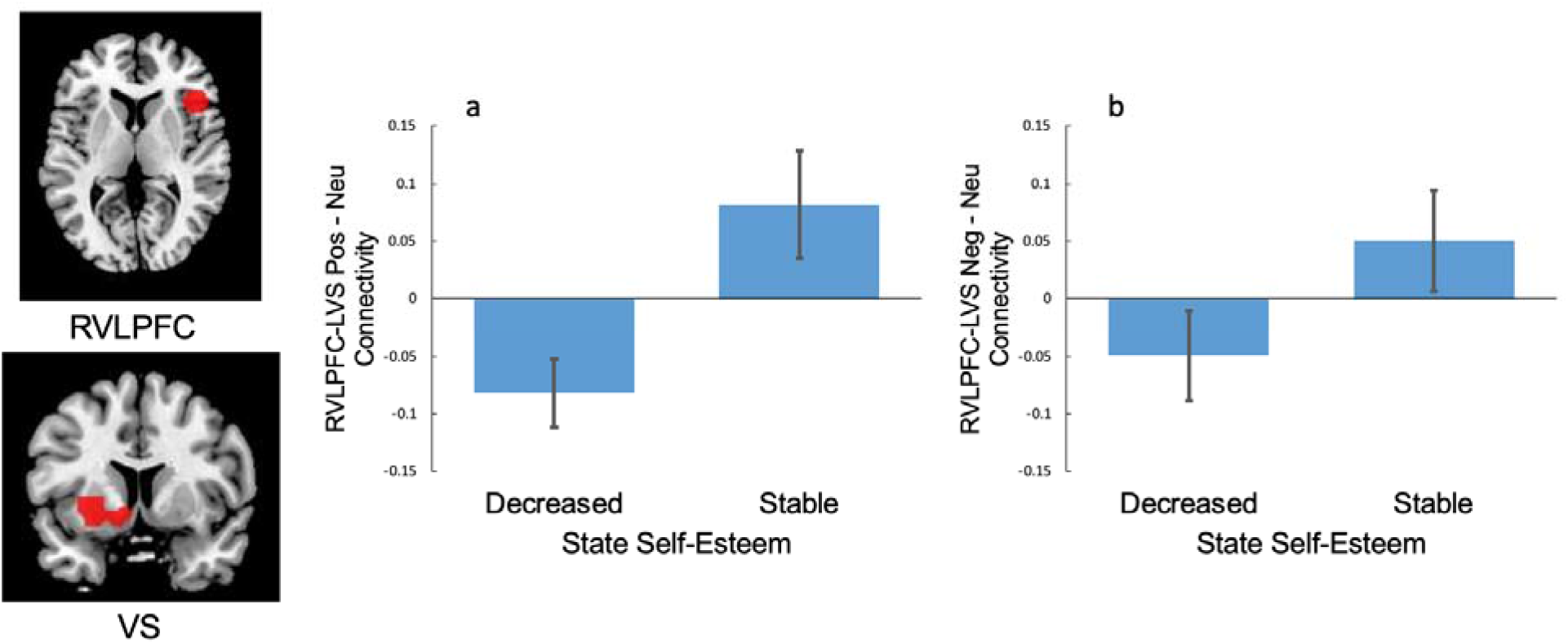
Bar graph depicting difference in RVLPFC-LVS functional connectivity during positive vs. neutral (left, a.) and negative vs. neutral social feedback (right, b.) between stable and decreased state self-esteem groups.

We next examined neural activity to negative (vs. neutral) feedback. Similar to what was observed in response to positive feedback, the stable state self-esteem group, relative to the decreased self-esteem group, showed significantly greater positive RVLPFC-LVS connectivity during negative (vs. neutral) feedback (*t*(37) = 1.70, *p* =.049; *U* = 130.00, *p* = .046, Fig. 1b). Results also indicated that those whose state self-esteem remained stable showed positive functional connectivity between these regions during negative (vs. neutral) feedback, whereas those whose state self-esteem declined showed average negative functional connectivity. Again, no differences between groups were found for functional connectivity between the RVLPFC and the right VS (*t*(37) = 0.59, *p* > .05; *U* = 219.00, *p* > .05).

#### Relationships with trait self-esteem

We next examined the relationship between trait self-esteem and RVLPFC-VS connectivity in response to both positive and negative feedback across the entire sample. Here, we found that, in response to positive (vs. neutral) feedback, trait selfesteem significantly correlated with RVLPFC-LVS connectivity (*r*(48) = .257, *p* = .036; Fig 2), such that individuals with higher trait self-esteem showed greater positive connectivity in the positive feedback condition relative to the neutral condition. These results remained the same when using a Spearman’s rank correlation analysis (*ρ*(48) = .238, *p* = .048). However, this same significant relationship did not hold for RVLPFC connectivity with the right VS (*r*(48) = .181, *p* = .104; *ρ*(48) = .187, *p* = .097). Finally, when examining functional connectivity during negative (vs. neutral) feedback, trait self-esteem was not significantly positively correlated with either RVLPFC-LVS connectivity (*r*(48) = .120, *p* > .05; *ρ*(48) = .131, *p* > .05) or RVLPFC-RVS functional connectivity (*r*(48) = .143, *p* > .05; *ρ*(48) = .084, *p* = .282).

**Figure 2.**
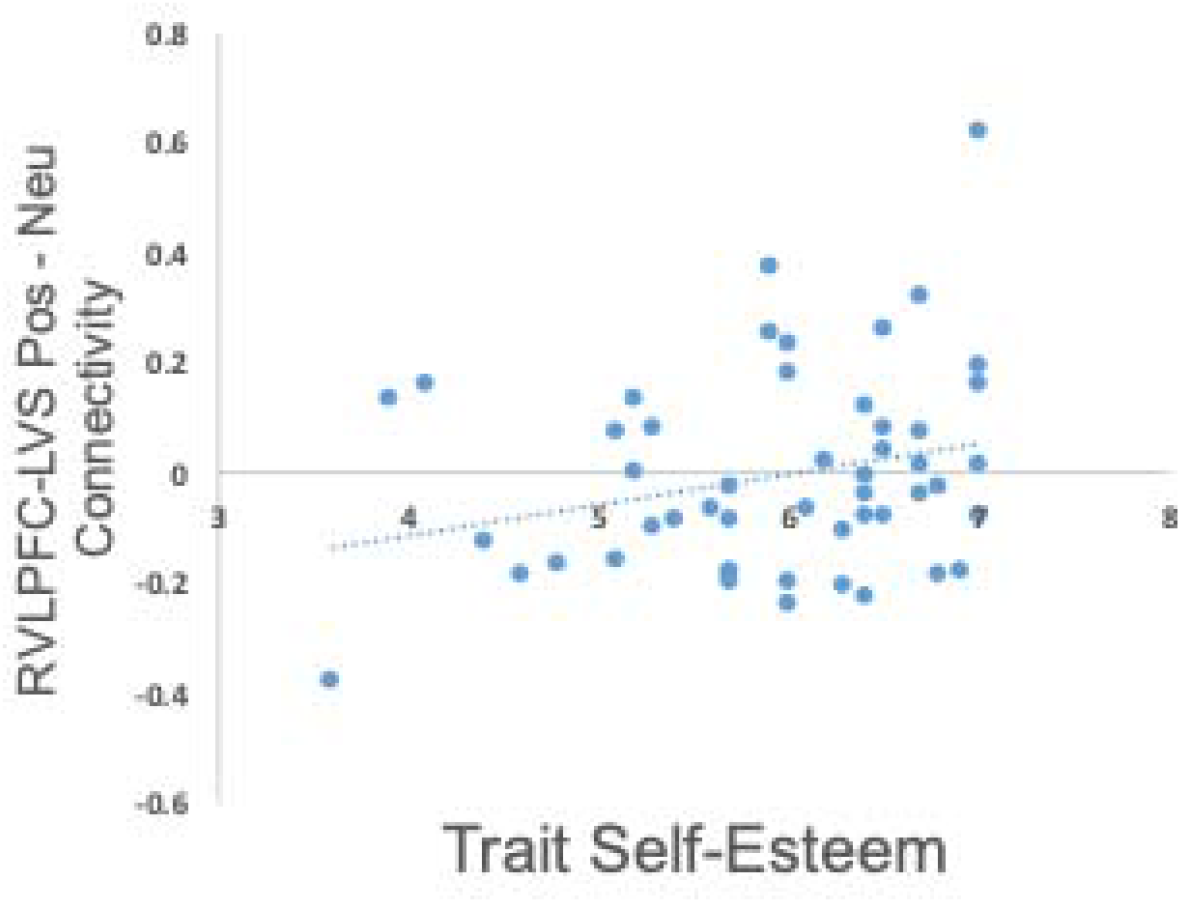
Scatterplot depicting the significant relationship between trait self-esteem and RVLPFC-LVS functional connectivity during positive vs. neutral social feedback.

## Discussion

The goal of the present research was to examine whether measures of self-enhancement were related to functioning within neural circuitry previously linked to emotion regulation, particularly regulatory processes implicated in increasing positive affect. In particular, we investigated whether frontostriatal functional connectivity during a social evaluation task positively related to changes in state self-esteem in response to socially evaluative feedback as well as individual differences in trait self-esteem. The current research builds upon earlier findings by examining self-enhancing tendencies with a functional connectivity approach, using a more naturalistic behavioral task, and by targeting a previously unexamined measure of neural circuit functioning linked to emotion regulation, namely connectivity between the RVLPFC and the VS.

Results indicated that individuals with greater positive connectivity between the RVLFPC and LVS during positive as well as negative feedback showed more stable state selfesteem compared to those whose self-esteem decreased in response to the evaluative feedback. In addition, greater positive connectivity between the RVLFPC and LVS during positive (vs. neutral) feedback was associated with relatively higher levels of trait self-esteem. These findings are in line with research demonstrating the importance of self-enhancement as a coping mechanism which may serve to upregulate or restore positive affect in reaction to threatening situations. Moreover, this pattern of results highlights the need to investigate regulatory mechanisms underlying the maintenance of self-esteem involved during not only negative emotion processing, but also positive emotion processing.

To date, the vast majority of social and affective neuroscience research on emotion regulation has focused on downregulating negative emotions. The current set of results point to the possibility that self-enhancement may be important, not only for decreasing negative affect, but for increasing positive affect as well, in order to restore feeling of self-worth in response to critical social evaluation. The present results which emphasize the importance of functional connectivity between RVLPFC and VS are in line with evidence showing that these regions are important for regulating positive affect. For example, both upregulation and downregulation of positive affect has been linked to ventral striatum activity (Greening et al., 2014; Kim & Hamann, 2007; Wager et al., 2008). Critically, RVLPFC activity has also been shown to directly correlate with subjective ratings of increased positive affect during emotion regulation tasks (Kim & Hamann, 2007). This is notable given that most emotion regulation research has been focused on the role of the RVLPFC in regulating negative affect. The current study highlights the RVLPFC-VS circuitry as another important neural pathway involved in the regulation of emotional well-being and self-worth.

In addition, the present results are well-aligned with social psychology behavioral research emphasizing the importance of upregulating positive affect, for the maintenance of self-enhancing tendencies. For example, behavioral research suggests individuals maintain positive self-views by engaging regulatory processes in order to “savor” or enhance positive feelings in the context of self-relevant events (Bryant, 1989, 2003; Wood et al., 2003, 2005). Notably, self-enhancing individuals have been shown to enjoy and magnify their success (Wood et al., 2003), whereas individuals with less self-enhancing tendencies fail to amplify the positive feelings associated with the event. Taken together, separate lines of previous research in both social psychology and neuroscience have both emphasized the importance of positive emotion regulatory processes for supporting well-being and self-worth; however, the current study is novel in its approach by demonstrating the importance of these processes for self-enhancement in a more naturalistic setting in which emotion regulation was not explicitly prompted.

While prior neuroimaging investigations into self-enhancement processes have been informative, many previous studies have also not utilized experimental paradigms simulating naturalistic social evaluative situations (c.f., Hughes & Beer, 2012b). This may be one of the reasons for previous lack of evidence linking the functioning of ventral striatum and its regulatory circuits to self-positivity biases and self-enhancing behaviors. For example, most investigations into the neural correlates of self-positivity biases, such as the “above average effect” (participants rating themselves more positively than statistically possible) have failed to reveal consistent activation in reward-related regions. Specifically, activation in the VS has shown no association with some measures while interestingly VMPFC has been shown to negatively correlate with self-enhancing behavior in some research (Beer & Hughes, 2010). Alternatively, other research has shown that measures of self-enhancement in reaction to specific threat conditions do indeed correlate with VMPFC activity (Hughes & Beer, 2012b). Hence, additional work is needed to further clarify the neural correlates of self-enhancement in response to self-threatening situations.

The current study may have multiple methodological advantages in comparison to prior studies. For one, the experimental paradigm leverages a naturalistic behavioral task with a high degree of psychological realism; that is, participants ostensibly believed that the social evaluative feedback was from similar peers and, as result, their levels of state self-esteem fluctuated as they would in real life situations. Relatedly, our functional connectivity approach allowed us to target regulatory processes in an unobtrusive manner, while not explicitly prompting conscious emotion regulation. These combined advantages allowed us to effectively examine the functioning of a putative positive emotional regulatory neural mechanism as it likely occurs in response to real world social evaluation.

Our pattern of neural findings, suggesting the importance of upregulating positive affect for self-worth, also reinforces the significance of similar findings in the clinical neuroscience of emotion regulation. Research shows that greater frontostriatal connectivity relates to greater self-reported positive affect among depressed patients, whereas a lack of frontostriatal connectivity is associated with depressive symptoms (Heller et al., 2013). Healthy profiles of functional connectivity between lateral PFC and VS have been shown to be sustained over time in healthy adults (Heller et al., 2009). However, depressed patients who are unable to sustain lateral PFC-striatum connectivity experience reduced positive affect. Research also shows that greater lateral PFC-VS connectivity is associated with lower levels of trait anhedonia in healthy adults (Keller et al., 2013). Importantly, tasks involving naturalistic mood induction link positive affect in healthy adults with frontostriatal connectivity (Admon & Pizzagalli, 2015). That we found frontostriatal connectivity to be related to self-enhancement seems quite reasonable in the light of the findings focused on clinical disorders, health and emotional well-being. Given our results, findings showing depressed patients have lower levels of frontostriatal functioning may also suggest that the inability to generate a sense of self-worth in depression is associated with connectivity in this circuit. Future research should explore this interesting potential direction.

While the pattern of observed results generally matched our prior hypotheses, a few noteworthy exceptions and study limitations are worth highlighting. First, although we predicted that trait self-esteem would be correlated with RVLPFC-VS functional connectivity in response to both positive (vs. neutral) and negative (vs. neutral) feedback, results showed trait self-esteem only correlated with connectivity in response to positive (vs. neutral) feedback. Although it’s not clear why, the fact that trait self-esteem did not correlate with connectivity to negative feedback could mean that trait self-esteem maintenance may have more to do with increasing positive affect (or savoring the experience) following positive experiences rather than increasing positive affect after negative experiences. It is also possible that the lack of effect could be due to being underpowered.

Second, we found hemispheric laterality effects in these results, such that the main set of significant associations were primarily found for the left VS. This is also somewhat unexpected given that there is a lack of definitive evidence for hemispheric specialization in VS functioning in the context of self-enhancement related processes. Regardless, prior studies targeting the RVLPFC-VS pathway to investigate the neural correlates of emotion regulation success have also found unexpected hemispheric differences. In particular, Wager et al. (2008) using a voxelwise mediation analysis found that the relationship between RVLPFC activity and emotion reappraisal success was specifically mediated by a cluster of activity within the left VS. It remains to be seen whether there will be definitive evidence for the specific involvement of RVLPFC-LVS circuit functioning in future neuroimaging studies on the topic of self-enhancement.

Lastly, while we believe that differences in RVLPFC-VS functional connectivity may be reflecting differences in positive emotion regulation in response to experimental stimuli, we lacked multiple convergent measures to definitively confirm this interpretation. Nonetheless, multiple studies which have investigated similar neural circuitry have found direct associations between frontostriatal functioning and positive affect (Admon & Pizzagalli, 2015; Heller et al., 2009; Keller et al., 2013). Moreover, previous researchers have also directly linked frontostriatal functioning to more active forms of explicit emotion regulation (Greening et al., 2014; Kim & Hamann, 2007; Wager et al., 2008). Further research is needed to definitively confirm associations between dynamic changes in self-worth, positive mood, and PFC-VS connectivity.

In summary, the results of the present study on self-enhancement inform current areas of research linking positive emotion regulation mechanisms (e.g., mechanisms involved in generating feelings of self-worth) to the functional interactions between the prefrontal cortex and striatum. Using a task simulating naturalistic social interaction, the current study showed that neural circuits connecting regulatory-related PFC regions and reward-related VS regions are likely involved in generating feelings of self-worth in response to threatening or challenging social feedback. Our findings support multiple lines of evidence demonstrating that the self-enhancement motive is underpinned by emotional regulation mechanisms which upregulate or restore positive affect in socially evaluative contexts. These results may have implications for clinical disorders such as depression, given that these disorders have been consistently linked with reduced functioning within frontostriatal circuits and are characterized by impaired selfesteem. Lastly, these findings underscore the importance of examining social processes with multiple types of functional neuroimaging measures (e.g., functional connectivity) and with paradigms that more closely approximate the real-world situations in which these social processes emerge.

## Notes

### Competing Interest Statement

The authors have declared no competing interest.

## References

Admon, R., & Pizzagalli, D. A. (2015). Corticostriatal pathways contribute to the natural time course of positive mood. Nature Communications, 6(1), 1–11. https://doi.org/10.1038/ncomms10065

Alicke, M. D. (1985). Global self-evaluation as determined by the desirability and controllability of trait adjectives. Journal of Personality and Social Psychology, 49(6), 1621–1630. https://doi.org/10.1037/0022-3514.49.6.1621

Alicke, M. D., & Sedikides, C. (2009). Self-enhancement and self-protection: What they are and what they do. European Review of Social Psychology, 20, 1–48. https://doi.org/10.1080/10463280802613866

Alicke, M. D., & Sedikides, C. (Eds.). (2011). Handbook of self-enhancement and self-protection. Guilford Press.

Beer, J. S. (2014). Exaggerated positivity in self□evaluation: A social neuroscience approach to reconciling the role of self□esteem protection and cognitive bias. Social and Personality Psychology Compass, 8(10), 583–594.

Beer, J. S., & Hughes, B. L. (2010). Neural systems of social comparison and the “above-average” effect. NeuroImage, 49(3), 2671–2679. https://doi.org/10.1016/j.neuroimage.2009.10.075

Beer, J. S., & Hughes, B. L. (2011). Self-enhancement: A social neuroscience perspective. Handbook of Self-Enhancement and Self-Protection, 49–65.

Bryant, F. B. (1989). A Four-Factor Model of Perceived Control: Avoiding, Coping, Obtaining, and Savoring. Journal of Personality, 57(4), 773–797. https://doi.org/10.1111/j.1467-6494.1989.tb00494.x

Bryant, F. B. (2003). Savoring beliefs inventory (SBI): A scale for measuring beliefs about savouring. Journal of Mental Health, 12(2), 175–196. https://doi.org/10.1080/0963823031000103489

Chavez, R. S., & Heatherton, T. F. (2015). Multimodal frontostriatal connectivity underlies individual differences in self-esteem. Social Cognitive and Affective Neuroscience, 10(3), 364–370. https://doi.org/10.1093/scan/nsu063

Chavez, R. S., Heatherton, T. F., & Wagner, D. D. (2017). Neural population decoding reveals the intrinsic positivity of the self. Cerebral Cortex, 27(11), 5222–5229.

Eisenberger, N. I., Inagaki, T. K., Muscatell, K. A., Byrne Haltom, K. E., & Leary, M. R. (2011). The neural sociometer: Brain mechanisms underlying state self-esteem. Journal of Cognitive Neuroscience, 23(11), 3448–3455. https://doi.org/10.1162/jocn_a_00027

Fiske, S. T. (2019). Social beings: Core motives in social psychology (4th edition). John Wiley & Sons, Inc.

Flagan, T., Mumford, J. A., & Beer, J. S. (2017). How do you see me? The neural basis of motivated meta-perception. Journal of Cognitive Neuroscience, 29(11), 1908–1917.

Frank, D. W., Dewitt, M., Hudgens-Haney, M., Schaeffer, D. J., Ball, B. H., Schwarz, N. F., Hussein, A. A., Smart, L. M., & Sabatinelli, D. (2014). Emotion regulation: Quantitative meta-analysis of functional activation and deactivation. Neuroscience & Biobehavioral Reviews, 45, 202–211.

Greening, S. G., Osuch, E. A., Williamson, P. C., & Mitchell, D. G. V. (2014). The neural correlates of regulating positive and negative emotions in medication-free major depression. Social Cognitive and Affective Neuroscience, 9(5), 628–637. https://doi.org/10.1093/scan/nst027

Heimpel, S. A., Wood, J. V., Marshall, M. A., & Brown, J. D. (2002). Do people with low self-esteem really want to feel better? Self-esteem differences in motivation to repair negative moods. Journal of Personality and Social Psychology, 82(1), 128–147. https://doi.org/10.1037//0022-3514.82.1.128

Heller, A. S., Johnstone, T., Light, S. N., Peterson, M. J., Kolden, G. G., Kalin, N. H., & Davidson, R. J. (2013). Relationships Between Changes in Sustained Fronto-Striatal Connectivity and Positive Affect in Major Depression Resulting From Antidepressant Treatment. American Journal of Psychiatry, 170(2), 197–206. https://doi.org/10.1176/appi.ajp.2012.12010014

Heller, A. S., Johnstone, T., Shackman, A. J., Light, S. N., Peterson, M. J., Kolden, G. G., Kalin, N. H., & Davidson, R. J. (2009). Reduced capacity to sustain positive emotion in major depression reflects diminished maintenance of fronto-striatal brain activation. Proceedings of the National Academy of Sciences, 106(52), 22445–22450. https://doi.org/10.1073/pnas.0910651106

Hughes, B. L., & Beer, J. S. (2012a). Medial orbitofrontal cortex is associated with shifting decision thresholds in self-serving cognition. NeuroImage, 61(4), 889–898. https://doi.org/10.1016/j.neuroimage.2012.03.011

Hughes, B. L., & Beer, J. S. (2012b). Protecting the Self: The Effect of Social-evaluative Threat on Neural Representations of Self. Journal of Cognitive Neuroscience, 25(4), 613–622. https://doi.org/10.1162/jocn_a_00343

Izuma, K., Kennedy, K., Fitzjohn, A., Sedikides, C., & Shibata, K. (2018). Neural activity in the reward-related brain regions predicts implicit self-esteem: A novel validity test of psychological measures using neuroimaging. Journal of Personality and Social Psychology, 114(3), 343.

Keller, J., Young, C. B., Kelley, E., Prater, K., Levitin, D. J., & Menon, V. (2013). Trait anhedonia is associated with reduced reactivity and connectivity of mesolimbic and paralimbic reward pathways. Journal of Psychiatric Research, 47(10), 1319–1328. https://doi.org/10.1016/j.jpsychires.2013.05.015

Kim, S. H., & Hamann, S. (2007). Neural Correlates of Positive and Negative Emotion Regulation. Journal of Cognitive Neuroscience, 19(5), 776–798. https://doi.org/10.1162/jocn.2007.19.5.776

Kuzmanovic, B., Jefferson, A., & Vogeley, K. (2016). The role of the neural reward circuitry in self-referential optimistic belief updates. NeuroImage, 133, 151–162.

Leary, M. R., Haupt, A. L., Strausser, K. S., & Chokel, J. T. (1998). Calibrating the sociometer: The relationship between interpersonal appraisals and state self-esteem. Journal of Personality and Social Psychology, 74(5), 1290–1299. https://doi.org/10.1037//0022-3514.74.5.1290

Mezulis, A. H., Abramson, L. Y., Hyde, J. S., & Hankin, B. L. (2004). Is there a universal positivity bias in attributions? A meta-analytic review of individual, developmental, and cultural differences in the self-serving attributional bias. Psychological Bulletin, 130(5), 711–747. https://doi.org/10.1037/0033-2909.130.5.711

Moieni, M., Irwin, M. R., Jevtic, I., Breen, E. C., & Eisenberger, N. I. (2015). Inflammation impairs social cognitive processing: A randomized controlled trial of endotoxin. Brain, Behavior, and Immunity, 48, 132–138. https://doi.org/10.1016/j.bbi.2015.03.002

Muscatell, K. A., Eisenberger, N. I., Dutcher, J. M., Cole, S. W., & Bower, J. E. (2016). Links between inflammation, amygdala reactivity, and social support in breast cancer survivors. Brain, Behavior, and Immunity, 53, 34–38. https://doi.org/10.1016/j.bbi.2015.09.008

Preuss, G. S., & Alicke, M. D. (2009). Everybody loves me: Self-evaluations and metaperceptions of dating popularity. Personality & Social Psychology Bulletin, 35(7), 937–950. https://doi.org/10.1177/0146167209335298

Roese, N. J., & Olson, J. M. (2016). Better, Stronger, Faster: Self-Serving Judgment, Affect Regulation, and the Optimal Vigilance Hypothesis: *Perspectives on Psychological Science*. https://journals.sagepub.com/doi/10.1111/j.1745-6916.2007.00033.x

Rosenberg, M. (1965). Rosenberg self-esteem scale (RSE). Acceptance and Commitment Therapy. Measures Package, 61(52), 18.

Sedikides, C., Gaertner, L., & Toguchi, Y. (2003). Pancultural self-enhancement. Journal of Personality and Social Psychology, 84(1), 60–79. https://doi.org/10.1037/0022-3514.84.1.60

Sedikides, C., & Gregg, A. P. (2008). Self-Enhancement: Food for Thought: *Perspectives on Psychological Science*. https://journals.sagepub.com/doi/10.1111/j.1745-6916.2008.00068.x

Taylor, S. E., & Armor, D. A. (1996). Positive illusions and coping with adversity. Journal of Personality, 64(4), 873–898. https://doi.org/10.1111/j.1467-6494.1996.tb00947.x

Taylor, Shelley E., & Sherman, D. K. (2008). Self-enhancement and self-affirmation: The consequences of positive self-thoughts for motivation and health. In Handbook of motivation science (pp. 57–70). The Guilford Press.

Tesser, A. (2000). On the Confluence of Self-Esteem Maintenance Mechanisms. Personality and Social Psychology Review, 4(4), 290–299. https://doi.org/10.1207/S15327957PSPR0404_1

Tzourio-Mazoyer, N., Landeau, B., Papathanassiou, D., Crivello, F., Etard, O., Delcroix, N., Mazoyer, B., & Joliot, M. (2002). Automated anatomical labeling of activations in SPM using a macroscopic anatomical parcellation of the MNI MRI single-subject brain. Neuroimage, 15(1), 273–289.

Wager, T. D., Davidson, M. L., Hughes, B. L., Lindquist, M. A., & Ochsner, K. N. (2008). Prefrontal-subcortical pathways mediating successful emotion regulation. Neuron, 59(6), 1037–1050. https://doi.org/10.1016/j.neuron.2008.09.006

Wood, J. V., Heimpel, S. A., & Michela, J. L. (2003). Savoring Versus Dampening: Self-Esteem Differences in Regulating Positive Affect. Journal of Personality and Social Psychology, 85(3), 566–580. https://doi.org/10.1037/0022-3514.85.3.566

Wood, J. V., Heimpel, S. A., Newby-Clark, I. R., & Ross, M. (2005). Snatching Defeat From the Jaws of Victory: Self-Esteem Differences in the Experience and Anticipation of Success. Journal of Personality and Social Psychology, 89(5), 764–780. https://doi.org/10.1037/0022-3514.89.5.764

